## Supplemental Tables for "Two Distinct Regulatory Systems Control Pulcherrimin Biosynthesis in *Bacillus subtilis*"

**S1 Table.** Strains Used in this Study.

| Strain | Genotype | Selection |
| --- | --- | --- |
| DK1042 | *B. subtilis* NCBI 3610 ComI$^{Q12L}$ | - |
| NF057 | *scoC::erm* | MLS |
| NF062 | *yvmC::kan* | Kan |
| NF064 | *scoC::erm abrB::kan* | MLS Kan |
| NF077 | *abrB::kan* | Kan |
| NF079 | Δ*pchR abrB::kan* | Kan |
| NF081 | Δ*pchR* | - |
| NF088 | ΔpchR scoC::erm | MLS |
| NF089 | *scoC::erm yvmC::kan* | MLS Kan |
| NF092 | Δ*pchR scoC::cam abrB::erm* | Cam Erm |
| NF093 | *scoC::erm lacA::pscoC-scoC* | Cam Erm |
| NF102 | WT *amyE::pYvmC-GFP* | Cam |
| NF103 | *scoC::erm amyE::pYvmC-GFP* | Cam Erm |
| NF104 | Δ*pchR amyE::pYvmC-GFP* | Cam |
| NF105 | Δ*pchR scoC::erm amyE::pYvmC-GFP* | Cam Erm |
| NF117 | *abrB::kan amyE::pYvmC-GFP* | Kan Cam |
| NF106 | Δ*pchR abrB::kan amyE::pYvmC-GFP* | Kan Cam |
| NF107 | Δ*pchR* Δ*scoC abrB::erm amyE::pYvmC-GFP* | Erm Cam |

**S2 Table.** DNA fragments and Plasmids Used in this Study.

| Name | Description | Resistance |
| --- | --- | --- |
| fNLF001 | abrB::erm with flanking homologous regions | Erm |
| fNLF003 | pchR::erm with flanking homologous regions | Erm |
| fNLF005 | yvmC::kan with flanking homologous regions | Kan |
| fNLF011 | yvmC::erm with flanking homologous regions | Erm |
| fNLF015 | scoC::camR-pscoC-FLAG-ScoC with flanking homologous regions | Cam |
| fNLF016 | scoC::camR with flanking homologous regions | Cam |
| pNF035 | pYvmC in pDR110 | Spec |
| pNF038 | lacA::pscoC-ScoC | Cam |
| pNF039 | PchR in pE-SUMO | Kan |
| pNF040 | AbrB in pE-SUMO | Kan |
| pTMN007[#] | ScoC in pE-SUMO | Kan |
| pNF047 | pYvmC in pGFP-Star | Cam |

### - see [25].

**S3 Table.** Oligonucleotides Used in this Study.

| Name | Primer Description/Use | Sequence (5′-3′) | Amplicon Description |
|---|---|---|---|
| oNLF336 | abrB_KO_US_For | atagtatttcagaagacgatccgc | AbrB::AbR disruption |
| oNLF337 | abrB_KO_US_Rev | ctctcctttctcgcctgccattctcctcccaagagata | |
| oNLF338 | abrB_KO_DS_For | gcagtgacaggagcctcgtaatcatttcttgtacaaaa | |
| oNLF339 | abrB_KO_DS_Rev | aatgtaaggacaatagctggtatgc | |
| oNLF340 | pchR_KO_US_For | tactgatcttctcacccagcttc | PchR::AbR disruption |
| oNLF341 | pchR_KO_US_Rev | ctctcctttctcgcctgccataggctaccttctttctt | |
| oNLF342 | pchR_KO_DS_For | gcagtgacaggagcctcgtaaacaaaaggcggtgtac | |
| oNLF343 | pchR_KO_DS_Rev | cattgtgagcaagtaggcagatac | |
| oNLF344 | yvmC_KO_US_For | tcggtttgttctgcttcaagt | YvmC::AbR disruption |
| oNLF345 | yvmC_KO_US_Rev | ctctcctttctcgcctgccatctcattcacccctaaaa | |
| oNLF346 | yvmC_KO_DS_For | gcagtgacaggagcctcgtgataggggagtaaaacat | |
| oNLF347 | yvmC_KO_DS_Rev | gtcaggctgttcaaatgcttc | |
| oNLF356 | AbR_KO_For | gcaggcgagaaaggagag | AbR Amplification |
| oNLF357 | AbR_KO_Rev | cgaggctcctgtcactgc | |
| oNLF432 | pNF035_For_FAM | gctgcaggaattcgactctc | 5' FAM and 5' IRD700 probe (WT and Δ59) for DNAse I footprinting and EMSA |
| oNLF433 | pNF035_For_IRD700 | gctgcaggaattcgactctc | |
| oNLF387 | yvmC_promoter_reverse_pNF035 | catgtttgtcctccttattagttaatcagctagctccggtcatctcattcacc | |
| oLVG025A | Amplifies pDR110 backbone, with oNLF467 | ctcttgccagtcacgttacg | |
| oNLF467 | yvmC_Δ59_us_rev | taataatcattttcaccaaacgtcaatatgatctgtg | |
| oNLF468 | yvmC_Δ59_ds_for | catattgacgtttggtgaaaatgattattaaaatcttaaaaaacatttg | |
| oNLF407 | scoC_us_fwd | ctatcgtcagctttattgatc | scoC::cam disruption |
| oNLF408 | scoC_us_rev | cgctctcctttctcgcctgccattacgtcacctgcttc | |
| oNLF409 | camR_fwd | gcaggcgagaaaggagagcgtcaggtggcacttttcg | |
| oNLF410 | camR_rev | cgaggctcctgtcactgcgacattagaaaaccgactgtaaaaag | |

**S3 Table Cont.** Oligonucleotides Used in this Study.

| | | | |
|---|---|---|---|
| oNLF411 | scoC_ds_fwd | tcgcagtgacaggagcctcgaagagctcgaacctgtaaac | |
| oNLF412 | scoC_ds_rev | aacaagaatatccaagccg | |
| oNLF471 | lacA_US_For_pBR332 | gtaatgataccgatgaaacgagaggcgggacagatatcctcg | Native ScoC complementation at *lacA* |
| oNLF472 | lacA_US_Rev_pBR332 | gtgccacctgactcacattctcctccttgttc | |
| oNLF473 | CamR_for_lacA | aggagaatgtgagtcaggtggcacttttcg | |
| oNLF474 | pscoC-ScoC_rev_lacA | cggagcatcagcttaactgtttacaggttcg | |
| oNLF475 | lacA_ds_for_pBR332 | gtaaacagttaagctgatgctccgctcgatatg | |
| oNLF476 | lacA_ds_rev_pBR332 | acctacatctgtattaacgaagcggcctccattacatctcttactgc | |
| oNLF495 | abrB_pET-SUMO_for | attgaggctcaccgcgaacagattggaggtatgaaatctactggtattgtac | AbrB Purification Vector |
| oNLF496 | abrB_pET-SUMO_rev | gatctcagtggtggtggtggtggtgctcgattatttaaggttttgaagctg | |
| oNLF497 | pchR_pET-SUMO_for | attgaggctcaccgcgaacagattggaggtatgtctgatttgacaaaacag | PchR Purification Vector |
| oNLF498 | pchR_pET-SUMO_rev | gatctcagtggtggtggtggtggtgctcgattactttacaggtttgtctg | |
| oNLF524 | yvmC_upstream_for | ttacagtcattttaccgcgggctttcccatgtgatgtttacatttattttcaaatatttg | pyvmC-GFP integration vector |
| oNLF525 | yvmC_upstream_rev | cgttaccattccggtcttatcccgctttaagtc | |
| oNLF526 | yvmC_downstream_for | ttaaagcgggataagaccggaatggtaacggaaag | |
| oNLF527 | yvmC_downstream_rev | ctattgaatccatagtagttcctccttccctaaattgagctttcgccc | |
